# Thrombocytopenia and splenic platelet directed immune responses after intravenous ChAdOx1 nCov-19 administration

**DOI:** 10.1101/2021.06.29.450356

**Authors:** Leo Nicolai, Alexander Leunig, Kami Pekayvaz, Afra Anjum, Eva Riedlinger, Luke Eivers, Marie-Louise Hoffknecht, Dario Rossaro, Raphael Escaig, Rainer Kaiser, Vivien Polewka, Anna Titova, Karsten Spiekermann, Matteo Iannacone, Konstantin Stark, Steffen Massberg

## Abstract

Vaccines against SARS-CoV-2 are based on a range of novel vaccine platforms, with adenovirus-based approaches (like ChAdOx1 nCov-19) being one of them. Recently a rare and novel complication of SARS-CoV-2 targeted adenovirus vaccines has emerged: thrombosis with thrombocytopenia syndrome (TTS). TTS is characterized by low platelet counts, clot formation at unusual anatomic sites and platelet-activating PF4-polyanion antibodies reminiscent of heparin-induced thrombocytopenia. Here, we employ *in vitro* and *in vivo* models to characterize the possible mechanisms of this platelet-targeted autoimmunity. We show that intravenous but not intramuscular injection of ChAdOx1 nCov-19 triggers platelet-adenovirus aggregate formation and platelet activation. After intravenous injection, these aggregates are phagocytosed by macrophages in the spleen and platelet remnants are found in the marginal zone and follicles. This is followed by a pronounced B-cell response with the emergence of circulating antibodies binding to platelets. Our work contributes to the understanding of TTS and highlights accidental intravenous injection as potential mechanism for post-vaccination TTS. Hence, safe intramuscular injection, with aspiration prior to injection, could be a potential preventive measure when administering adenovirus-based vaccines.

## Main text

The emergence of SARS-CoV-2 and the ensuing pandemic has led to the development of vaccines eliciting an immune response to the virus spike protein in an unprecedented time frame. Multiple vaccine platforms have been deployed successfully, with mRNA and adenovirus-vector based vaccines carrying the bulk of global vaccination efforts [1]. Recently, a potentially novel adverse effect of vaccination termed thrombosis with thrombocytopenia syndrome (TTS) or vaccine induced thrombotic thrombocytopenia (VITT) has been described in patients receiving adenovirus-based vaccines (ChAdOx1 nCov-19 (AstraZeneca) and Ad26.COV2.S (Johnson&Johnson) 4-30 days prior [2, 3]. This rare disease resembles heparin induced thrombocytopenia type II (HIT II) with positive PF4-polyanion antibody assay required for diagnosis [4, 5]. However, the underlying mechanism of TTS development only in a fraction of vaccinated individuals remain poorly understood and potential preventive measures are not established[6].

A previously healthy 27-year-old male presented to our emergency department with new onset headache non-responsive to ibuprofen 10 days after vaccination with ChAdOx1 nCov-19 (Fig. 1a). Initial evaluation revealed thrombocytopenia and elevated D-Dimer, as well as a positive rapid HIT screening test (Suppl. Fig. 1a and Suppl. Table 1 & 2). A contrast enhanced head CT scan was concordant with sinus vein thrombosis (SVT), an otherwise uncommon site of thrombosis suggestive of TTS (Fig. 1b). The patient was treated with IVIGs and anticoagulated with argatroban. He was discharged upon disappearance of symptoms and anticoagulation was continued with dabigatran. Thrombophilia screening was inconclusive (Suppl. Table 1). Evaluation of risk factors revealed current smoking. Tests used for the detection/confirmation of relevant PF4/polyanion antibodies (HIT-ELISA, HIT-IL-ACUSTAR and HIPA/PIPA test) were all negative in this patient, formally ruling out TTS according to guidelines[5] (Suppl. Table 2). Consistent with the case described here, reports on thrombotic and thrombocytopenic complications after adenovirus-based SARS-CoV-2 vaccination include an increased rate of isolated thrombocytopenia, as well as PF4-polyanion negative cases of sinus vein thromboses [7, 8]. We therefore asked if additional mechanisms beyond PF4/polyanion antibodies - potentially involving direct adenoviral vector-platelet interaction - might be involved in the pathogenesis of TTS.

**Figure 1.**
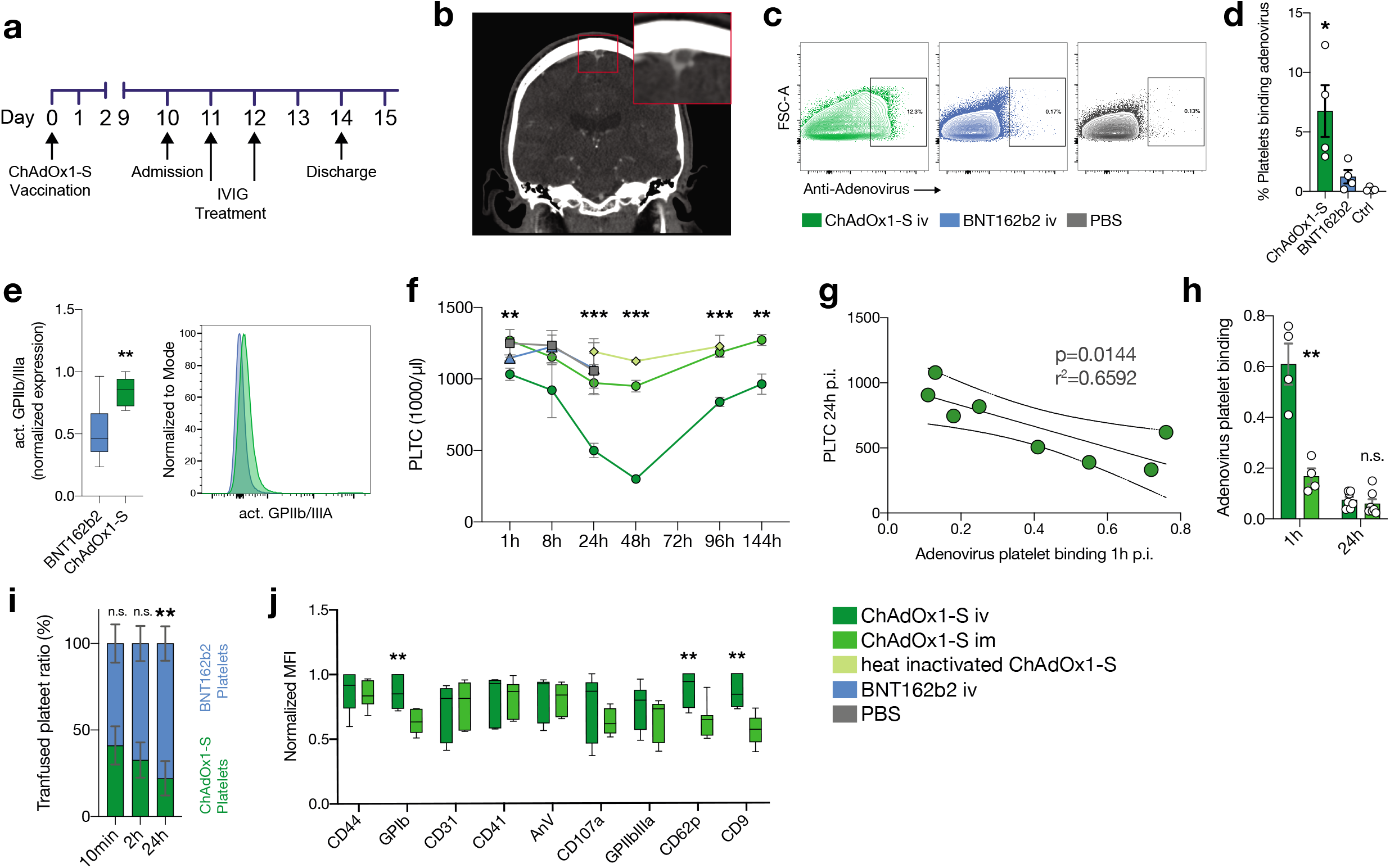
Intravenous but not intramuscular injection of ChAdOx1 nCov-19 triggers ChAdOx1 nCov-19-platelet binding and thrombocytopenia. **a**, Clinical time course of TTS patient, starting at vaccination with ChAdOx1-S (day 0). **b,** Coronal contrast enhanced computed tomography image of showing a likely sinus vein thrombosis (enlarged on the upper right). **c,** Exemplary flowcytometric contour plots of human washed-platelets with ChAdOx1 nCov-19, BNT162b2, or PBS. 2% contour with outliers shown gate shows adenovirus positive platelets. **d,** Quantification of adenovirus platelet binding according to **c.** One-way ANOVA with post-hoc Tukey’s test. Comparison of ChAdOx1 nCov-19 to both controls. n≥3 human donors per group. **e,** Platelet expression of activated GPIIbIIIa incubated with ChAdOx1 nCov-19 or BNT162b2. Normalized MFIs. Unpaired t-test. n=8 donors per group. Exemplary histogram of activated GPIIbIIIa is also shown. **f,** Platelet counts of mice over time. Multiple t-tests with Holm-Sidak correction of ChAdOx1 nCov-19 i.v. and i.m. is shown. n≥3 per time point for of ChAdOx1 nCov-19 groups, n≥2 per time point for other groups. **g,** Linear regression of platelet count at 24h p.i. and adenovirus binding to platelets 1h p.i.. 95% confidence interval shown, p value denotes line deviation from zero. **h,** Adenovirus-platelet binding after i.m. or i.v. administration of ChAdOx1 nCov-19 1h and 24h post inoculation. Unpaired t-tests. n≥4 per group. **i,** Ratio of transfused BNT162b2 and ChAdOx1 nCov-19 platelets (total transfused platelets are normalized to 100%) over time. Unpaired t-tests. n=4 per time point. **j,** Platelet surface marker expression of mouse platelets 24h after administration of ChAdOx1 nCov-19 i.v. or i.m.. Normalized MFIs. Multiple t-tests with Holm-Sidak correction. n=7 mice per group. Error bars are mean ±s.e.m. *p<0.05, **p<0.01, ***p<0.001.

To study direct adenoviral vector-platelet interaction, we incubated healthy human platelets with ChAdOx1 nCov-19, mRNA-based vaccine BNT162b2 (Pfizer-BioNTech) or phosphate buffered saline (PBS). Incubation with anti-pan-Adenovirus antibody revealed binding of ChAdOx1 nCov-19 to human platelets (Fig. 1c-d). This binding resulted in a shift of platelets towards an activated phenotype with increased activated GPIIb-IIIa integrin (Fig. 1e, Suppl Fig. 1b). Similarly, mouse platelets also strongly bound ChAdOx1 nCov-19 *in-vitro* (Suppl Fig. 1c).

Vaccines are routinely administered intramuscularly (i.m.) and trigger immune responses mainly in the draining lymph nodes [9]. Based on our finding that adenoviral vaccine binds to blood platelets, we hypothesized that accidental intravenous injection of adenoviral vaccine might lead to platelet-adenovirus aggregate formation with platelet activation and possibly platelet clearance. Indeed, intravenous but not i.m. injection of ChAdOx1 nCov-19 triggered transient thrombocytopenia in mice (Fig. 1f). Intravenous injection of PBS, BNT162b2 or heat inactivated ChAdOx1 nCov-19 injection had no effect on platelet counts (Fig. 1f). Thrombocytopenia induced by intravenous injection of ChAdOx1 nCov-19 was not due to fluid shift or changes in platelet production, as indicators like platelet distribution width, mean platelet volume and platelet larger cell ratio (P-LCR) did not show variations (Suppl Fig. 1d). The decline in platelet count was dose-dependent and correlated directly with adenovirus-positive platelets circulating in the blood one hour after i.v. injection (Fig. 1g, Suppl Fig. 1e). AID^-/-^ sIgM^-/-^ mice lacking IgM and IgG antibodies displayed the same platelet count kinetic, indicating that development of thrombocytopenia within 48hr after intravenous injection of ChAdOx1 nCov-19 does not depend on antibodies (Suppl. Fig. 1f). However, intravenous but not i.m. injection of ChAdOx1 nCov-19 resulted in a strong increase in platelet-adenovirus aggregates (Fig. 1h). These aggregates had a short dwell time and rapidly disappeared from the circulation (Fig. 1h). Disappearance of platelet-adenovirus aggregates preceded recovery of platelet counts (Fig 1f and h). To further investigate if binding of adenovirus is responsible for enhanced platelet clearance, we co-injected labeled platelets incubated with either ChAdOx1 nCov-19 or BNT162b2. We observed that ChAdOx1 nCov-19 incubated platelets were selectively removed from the circulation (Fig.1i) (2.8±1.1h half-life for ChAdOx1 nCov-19 incubated platelets vs. 10±3.7h half-life for BNT162b2 incubated platelets). At 24h after intravenous ChAdOx1 nCov-19 administration, mouse platelets displayed an activated phenotype with increased P-Selectin (CD62P), compared to intramuscular injection (Fig. 1j).

Platelets are known to bind and shuttle blood-borne pathogens to professional phagocytes in the spleen and liver to mount an adaptive immune response [10–12]. In the spleen, platelets are abundant in the red pulp during steady state conditions, but deposit pathogens in the marginal zone and follicles, which are predominant regions of adaptive immune responses, upon pathogen binding [13]. We therefore analyzed platelet localization after i.v. compared to i.m. application in the spleen. To robustly label all platelets, we injected a fluorescent non-activating Gp1b-antibody at the time of vaccine injection. Analysis of overall Gp1b content in the spleen after i.v. injection in contrast to i.m. injection revealed prominent accumulation of Gp1b+ material in the marginal zone and follicles; these areas are devoid of platelets under steady state conditions (Fig. 2a). In i.m injected animals, the Gp1b signal identified round and intact platelets in the splenic red pulp (Fig. 2a). In contrast, the Gp1b signal in the i.v. injection group appeared morphologically different with follicular and marginal zone clusters reminiscent of platelet remnants (Fig. 2a, Suppl. Fig. 1g). Professional phagocytes, particular red pulp macrophages, phagocytose activated or opsonized platelets in the spleen [14]. In line with this, we found increased platelet uptake by splenic F4/80 macrophages after ChAdOx1 nCov-19 i.v. compared to i.m. administration (Fig. 2b). In summary, these data show phagocytic uptake and processing of platelets and adenovirus in the spleen.

**Figure 2.**
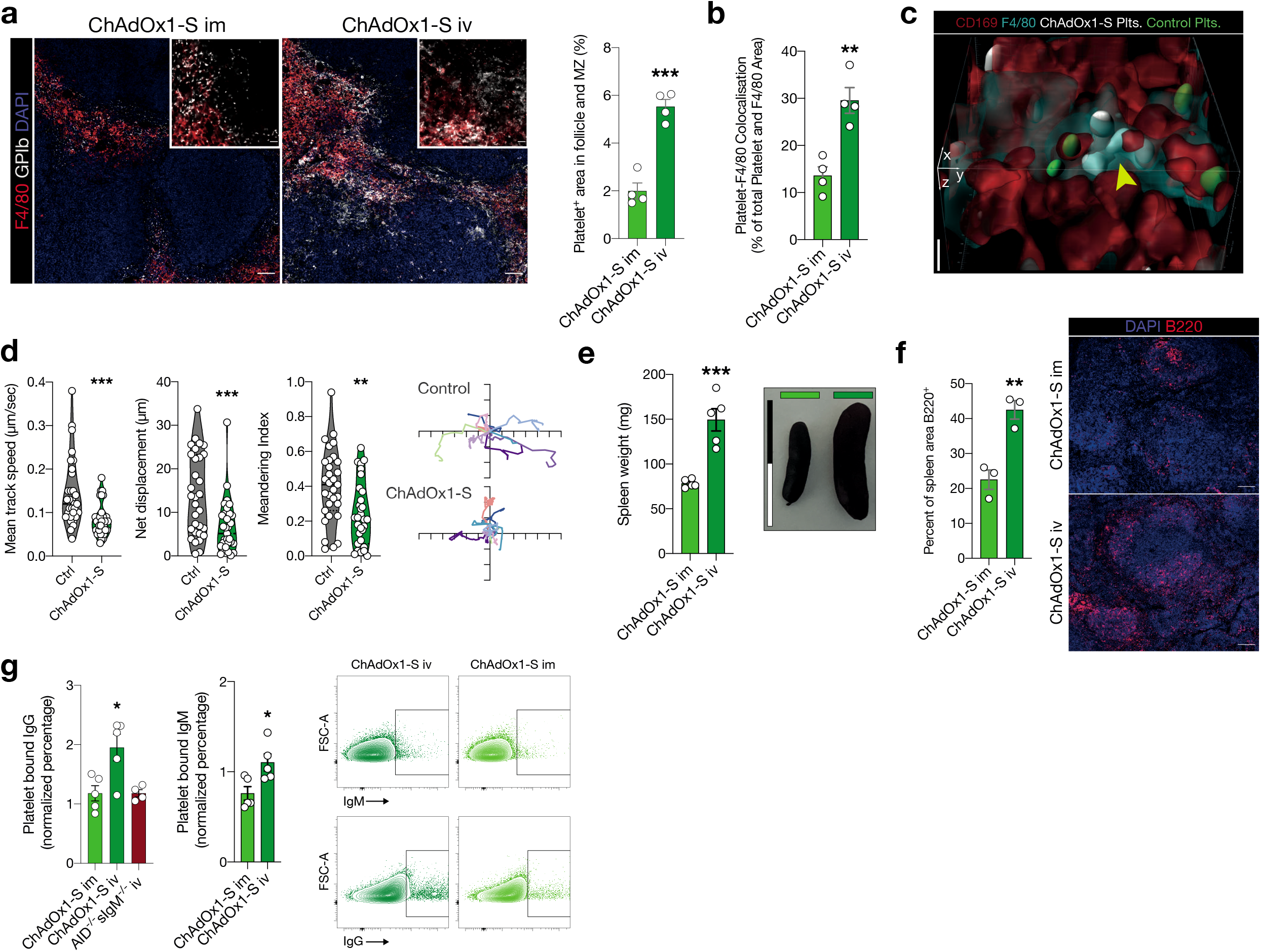
Platelet-adenovirus aggregates are taken up by macrophages in the spleen and lead to adenovirus-platelet directed immune responses. **a,** Images of X649 endogenous platelet labelling mice 24h after i.m. and i.v. ChAdOx1 nCov-19 administration. Cut out of red pulp to follicle transition is shown on the upper left. Scale bars for overview are 50μm, for cut outs 10μm. Quantification of Platelet (X649) area in the marginal and follicle zone as percent of marginal and follicular area. n=4 per group, unpaired t-test. **b,** Quantification of platelet-F4/80 co-localization in the splenic red pulp. Co-localization is shown as percent of total platelet-F4/80 area. n=4 per group, unpaired t-test. **c,** 3D reconstruction of in-vivo microscopy of the spleen with transfused platelets pretreated with either ChAdOx1 nCov-19 (white) or PBS (green). ChAdOx1 nCov-19 pretreated platelets are taken up by F4/80^+^/CD169^+^ macrophages, while control treated platelets are not. Scale bar is 5μm. **d,** Platelet tracking parameters of intraviral imaging. n=31 platelets tracked per group of one mouse. 10 exemplary tracked paths per group are shown on the left as x- and y-displacement, ticks signify 5μm intervals. Unpaired t-test with Welch’s correction. **e,** Spleen weights of animals 6d p.i.. Representative images of spleens of both groups are shown, scale bar is 2cm. Unpaired t-test. n=5 per group. **f,** Quantification of B220^+^ are in the spleen of i.v. or i.m. ChAdOx1 nCov-19 injected mice 6d p.i. as percentage of total splenic area. Unpaired t-test, n=3 per group. Representative images of a splenic micrograph of both groups are shown, scale bar is 100μm. **g,** Control mouse platelets positive for bound IgM and IgG after incubation with plasma from mice 6d p.i. with either i.m. or i.v. ChAdOx1 nCov-19 or ChAdOx1 nCov-19 AID^-/-^sIgM^-/-^ mice for IgG additionally. n=5 per group for i.v. or i.m., n=4 for AID^-/-^sIgM^-/-^. Unpaired t-tests between i.v. ChAdOx1 nCov-19 and the other groups. Percent of platelets binding IgM or IgG are normalized to the mean of 4 control plasmas. Error bars are mean ±s.e.m. *p<0.05, **p<0.01, ***p<0.001.

Even under steady-state conditions, contacts between passing platelets and local macrophages are frequently observed in the red pulp[15]. To understand the dynamics of platelet-adenovirus trafficking in more detail, we therefore co-transfused labeled control and ChAdOx1 nCov-19 incubated platelets and performed 4D intravital microscopy of the spleen (Suppl. Fig. 1h). We observed increased phagocytosis of ChAdOx1 nCov-19 pretreated platelets by CD169^+^ and F4/80^+^ macrophages in vivo (Fig 2c, Suppl Fig 1i-j), again pointing towards preferential targeting of virus-loaded platelets to professional phagocytes in the spleen. As a result, single-cell tracking in the spleen revealed decreased motility and enhanced adherence of the ChAdOx1 nCov-19 pretreated platelets to macrophages (Fig. 2d and Suppl. Video 1).

Based on the platelet uptake by splenic macrophages, we hypothesized that presentation of platelet antigens in the spleen might trigger anti-platelet antibody production resulting in a second, delayed wave of platelet depletion occurring with a characteristic lag time[16]. This resembles the kinetics of clinically apparent thrombocytopenia of TTS patients reported to occur 4-30 days post vaccination. To test this hypothesis, we assessed adaptive immune responses in the spleen upon i.m./i.v. ChAdOx1 nCov-19 application. Intravenous ChAdOx1 nCov-19 led to significant splenomegaly with enlarged B cell follicles in the spleen (Fig. 2e-f). Next, we determined if this immune response results in production of antibodies targeting platelets. We incubated control mouse platelets with plasma from i.m./i.v. ChAdOx1 nCov-19 treated animals 6 days post injection and control plasma. Naïve ChAdOx1 nCov-19 unexperienced platelets incubated with plasma from intravenously injected mice showed increased IgM and IgG binding, pointing towards circulating anti-platelet antibodies (Fig. 2g). These did not lead to platelet activation, most likely because mouse platelets unlike their human counterparts lack Fc-receptors (Suppl. Fig 1k). As expected, plasma from ChAdOx1 nCov-19 intravenously injected antibody deficient AID^-/-^ sIgM^-/-^ showed a signal comparable to controls. We also screened the patient who presented with PF4/polyanion antibody negative thrombocytopenia and SVT for antiplatelet antibodies. We found anti-platelet antibodies in this patient that were reactive to the platelet receptor GPIIb/IIIa, , although this result is limited by the fact that diagnostics were performed after IVIG treatment (Suppl Table 3).

Here, we use in vitro and murine models of ChAdOx1 nCov-19 vaccination to show that intravenous injection of this vaccine triggers platelet-adenovirus vector aggregate formation and platelet activation. This leads thrombocytopenia and uptake of these aggregates by professional phagocytes in the spleen. As a result, platelet and adenovirus antigens trigger the formation of platelet directed autoantibodies, as demonstrated by increased circulating anti-platelet IgG and IgM. Our findings might explain the rare incidence of TTS, as accidental intravenous deltoid injection is considered an unlikely event for anatomical reasons [17]. The CDC does not recommend aspiration during intramuscular application of vaccines [18]. However, in the case of adenovirus-based vaccines, aspiration prior to injection could offer a simple preventive measure against TTS.

So far, TTS has been linked exclusively to the presence of platelet-activating PF4-polyanion antibodies [3]. Here we report a patient with a TTS-like syndrome and no detectable PF4-polyanion antibodies similar to published reports [7, 8]. Non-platelet-activating PF4/polyanion antibodies appear only in low frequencies in both adenovirus- and mRNA-vaccinated patients, making it unlikely that antibody formation against these targets is common after vaccination [19, 20].

It is important to note that we used an animal model which inherently cannot fully recapitulate disease pathophysiology observed in humans. Murine platelets do not express Fcy receptors, and therefore are not suited as a model of antibody mediated platelet activation and thrombocytopenia, i.e. upon the injection of heparin and PF4/polyanion antibody [21]. Similarly, we did not observe delayed antibody-dependent thrombocytopenia after i.v. adenovirus injection in mice and did not detect platelet activation upon antibody binding.

Nevertheless, our data provide first experimental support for the potential sequence of events that could lead to TTS in some patients: Thrombocytopenia might be a result of accidental intravenous vaccine injection with ensuing platelet-adenovirus aggregation, which in turn triggers platelet activation and subsequent splenic clearance (Suppl. Fig. 3). This process is antibody independent and most likely clinically inapparent. Phagocytosis and processing of platelet-adenovirus aggregates by professional phagocytes then creates a cross-reactive immune response, which culminates in the delayed emergence of platelet-binding-antibodies. These platelet-binding-antibodies might induce a second wave of clinically apparent thrombocytopenia through platelet activation or opsonization. In fact, pro-thrombotic antibodies present in heparin induced thrombocytopenia Type II, and platelet opsonization with antibodies is a typical feature of immune thrombocytopenic purpura (ITP) Antibodies targeting platelets might shift these platelets towards a more thrombotic phenotype, which then leads to thrombosis in individuals with additional risk factors, including female sex, smoking or contraception. [22, 23]. Interestingly, we also detected platelet autoantibodies in the described patient case. Further research is needed to fully characterize the antibody responses in patients with TTS, as well as animal studies to define mechanisms and targets of the mounted adaptive immune response in depth.

## Suppl. Methods section

### Patient data

The patient was treated at the university hospital LMU Munich. The patient consented to taking part in this study. The study was approved by local ethics committee of Munich. Data on presentation, treatment and outcome are depicted in Fig 1a-b, Suppl. Fig 1a and the laboratory values table (Suppl Table 1-3). A HIT-IL-Acustar assay and a HYPHEN BioMed HIT-ELISA Test deemed sensitive for TTS returned negative for this patient [5]. PF4 induced platelet activation(PIPA) and heparin induced platelet activation (HIPA) tests were performed as previously reported[24].

**Suppl. Table 1.**
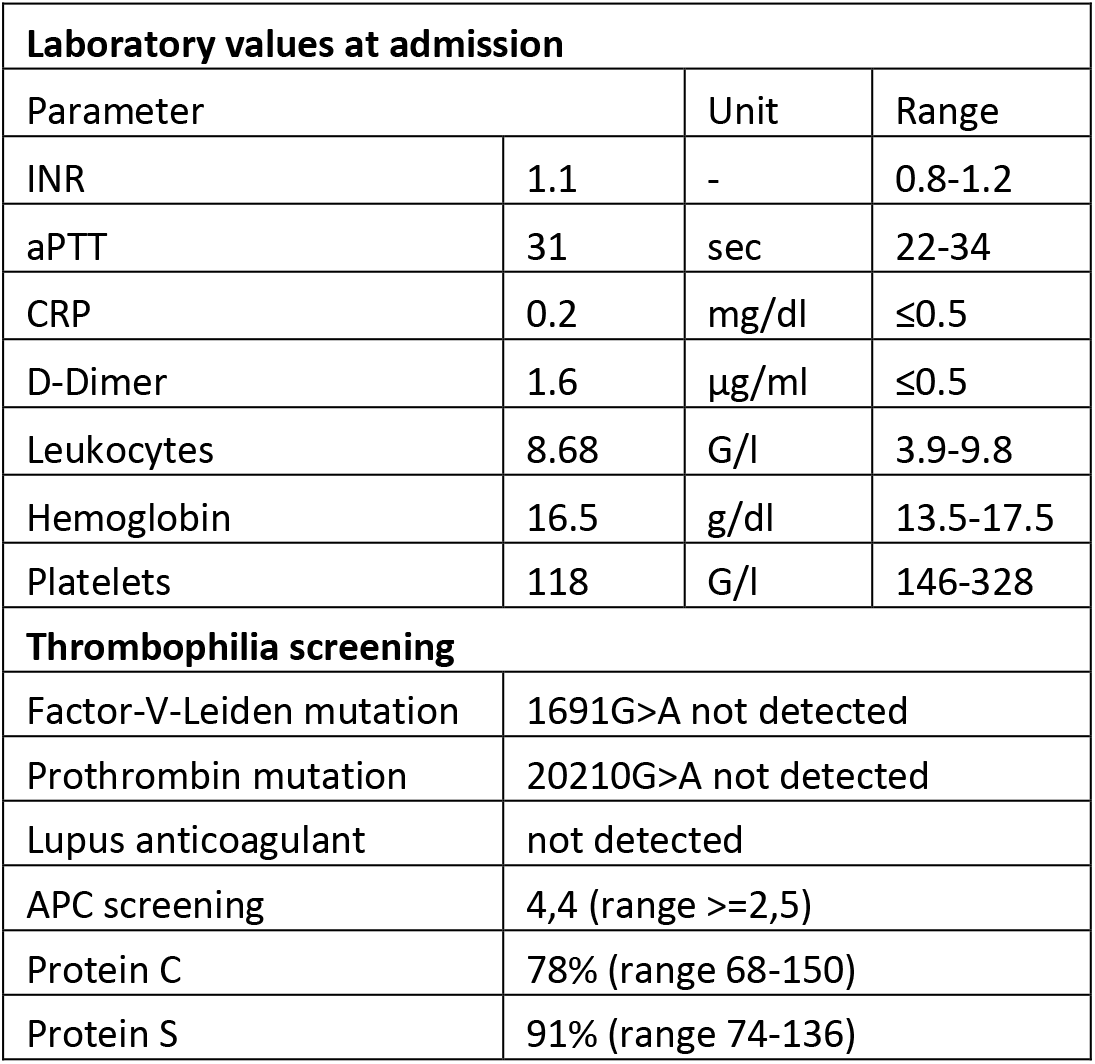

**Suppl. Table 2.**
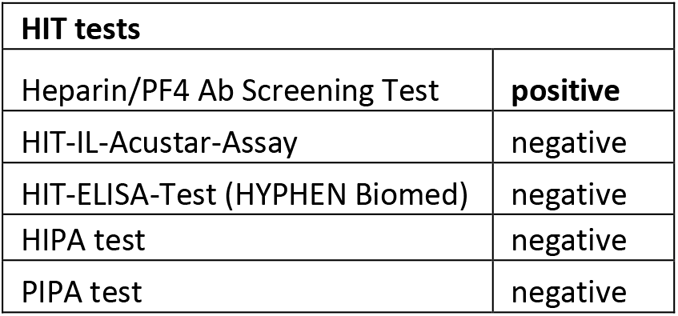

**Suppl. Table 3.**
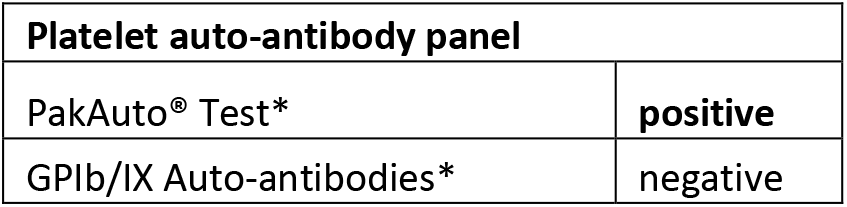
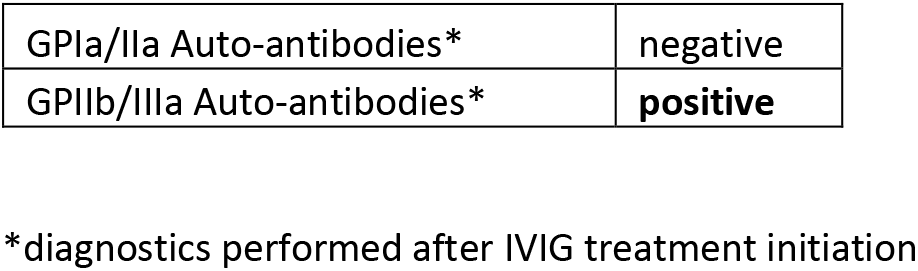

### Mouse Strains

C57BL/6 mice were purchased from The Jackson Laboratory. AID^-/-^ sIgM^-/-^ mice were bred and maintained at our animal facility. Mice were aged between 12-16 weeks for the experiments and were age and sex-matched for inter-group analyses. All mice live in standardized conditions where temperature, humidity, and hours of light and darkness are maintained at a constant level and provided water and food ad libitum. All animal experiments were performed in compliance with the relevant ethical regulations and were approved by the local administrative authority for the protection of animals (*Regierung von Oberbayern, München*).

### Vaccine acquisition

Only residual content of discarded BNT162b2 (Pfizer-BioNTech) and ChAdOx1 nCov-19 (AstraZeneca) vials, that, according to the supervising physicians’ body (*Kassenärztliche Vereinigung Bayerns*), could not be used for human application anymore, were used for this study. For ChAdOx1 nCov-19 this meant a storage period greater than 48h after first withdrawal of a dose from a multi-dose vial, for BNT162b2 it meant a storage period greater than 24h after first withdrawal of a dose from a multi-dose vial. Both vaccines were stored continuously at 4°C. Experiments were conducted as soon as possible after vaccine acquisition.

### Human platelet isolation

Human blood for *in-vitro* assays was obtained after informed consent from healthy male and female donors aged 20-35. Blood was collected with a syringe containing 1/7 of the blood volume citrate (39 mM citric acid, 75 mM sodium citrate, 135 mM dextrose; ACD). The blood was further diluted 1:1 with Tyrode’s buffer Tyrode’s buffer (137 mM NaCl, 2.8 mM KCl, 12 mM NaHCO3, 5.5 mM glucose, 10 mM HEPES, pH = 6.5), centrifuged at 70g for 35min, and the supernatant platelet-rich-plasma (PRP) was taken. PRP was further spun down after adding 0.1mg/ml PGI2 at 1200g for 10min. The resulting washed platelet pellet was resuspended in 7.2pH Tyrode’s buffer and used for subsequent experiments.

### Mouse platelet isolation

Mouse blood was obtained via retro-orbital blood collection, after anesthesia with medetomidine, midazolam and fentanyl (MMF). Blood was collected via a capillary and mixed with 1/7 of the blood volume citrate. The blood was further diluted 1:1 with Tyrode’s buffer (137 mM NaCl, 2.8 mM KCl, 12 mM NaHCO3, 5.5 mM glucose, 10 mM HEPES, pH = 6.5), centrifuged at 70g for 10min, and the supernatant platelet-rich-plasma (PRP) was taken. PRP was further spun at 1200g for 10min. The resulting washed platelet pellet was resuspended in 7.2pH Tyrode’s buffer and used for subsequent experiments.

### Platelet incubation with vaccines

Ca. 1×10^8^ washed platelets were incubated with ~5×10^7^ ChAdOx1 nCov-19 viral particles (100μl), or 100μl BNT162b2 or PBS for 20min. After this 1:200 X649 (mouse, emfret Analytics) or CD41 (human, Biolegend) was added and further incubated for 10min and then analyzed via flow-cytometry. For transfusion, after separate vaccine incubation with ChAdOx1 nCov-19 and BNT162b2, platelets were stained with 1:200 X649 or X488 (emfret Analytics) respectively for 10min. X488 and X649 labelling was switched for half of the mice to ensure no color bias. The platelets were spun down at 1200g for 10min, resuspended in 200μl 7.2pH Tyrode’s buffer and immediately injected separately via tail vein injection.

### Mouse vaccine injection and blood collection

5μl (low dose) or 50μl of ChAdOx1 nCov-19, BNT162b2, PBS, or heat inactivated ChAdOx1 nCov-19 were used for injections. Heat inactivation of ChAdOx1 nCov-19 was achieved by heating the vaccine to 99°C for 30min, then spinning the resulting protein precipitate down at 10,000g for 3min and taking the supernatant. Injection was performed either via tail vein or in the medial aspect of the thigh of isoflurane anesthetized mice. At specified time points, vaccine injected, and transfused mice were briefly anesthetized with isoflurane and blood was obtained via facial vein puncture and collected in EDTA capillaries. Blood counts were measured with the Sysmex XN-V Series XN-1000V cell counter, only blood counts that passed internal quality control were used.

### Platelet bound immunoglobulin detection

Mouse washed platelets were incubated with 1:4 plasma of mice 6d post-inoculation (p.i.) and control mice for 40min. After adding 1:100 CD41 PB, IgG Cy3 and IgM FITC and 20min incubation washed platelets were fixed, diluted and analyzed via flow cytometry.

### Intravital imaging of the spleen

Co-injection of platelets was performed as previously described[25]. After injection of X488 or X649 labelled platelets incubated with ChAdOx1 nCov-19 or PBS, mice were also injected with 15μl each of CD169 PE and F4/80 BV421 (Biolegend), anesthetized with MMF, shaven, and the spleen exposed via a small lateral cut. Subsequently the spleen was mobilized and imaged with a Zeiss LSM 880 confocal microscope (x20 obj.) in airyscan mode. Regular blood flow in the spleen was observed throughout imaging, and z-Stack images (1μm slice thickness) as well as videos were taken.

### Flow cytometry

Flow cytometry to characterize platelet profiles was performed as previously described [26]. Briefly, incubation with panels at 1:200 per antibody of either washed platelets or whole blood, at least 5x the amount of FACS Lysing Solution (BD) was added to fixate platelets and lyse blood. Flow cytometry was performed on a BD LSRFortessa and BD FACSCanto. Flowjo (BD) was used for flowcytometric analysis, gating strategies are shown in Supplemental Figure 2. For adenovirus binding, as well as IgM and IgG binding a positive gate was used, for MFIs the MFI of each marker per data point was normalized by dividing it by the highest MFI of that marker and experimental run.

Antibodies used for Flow Cytometry:

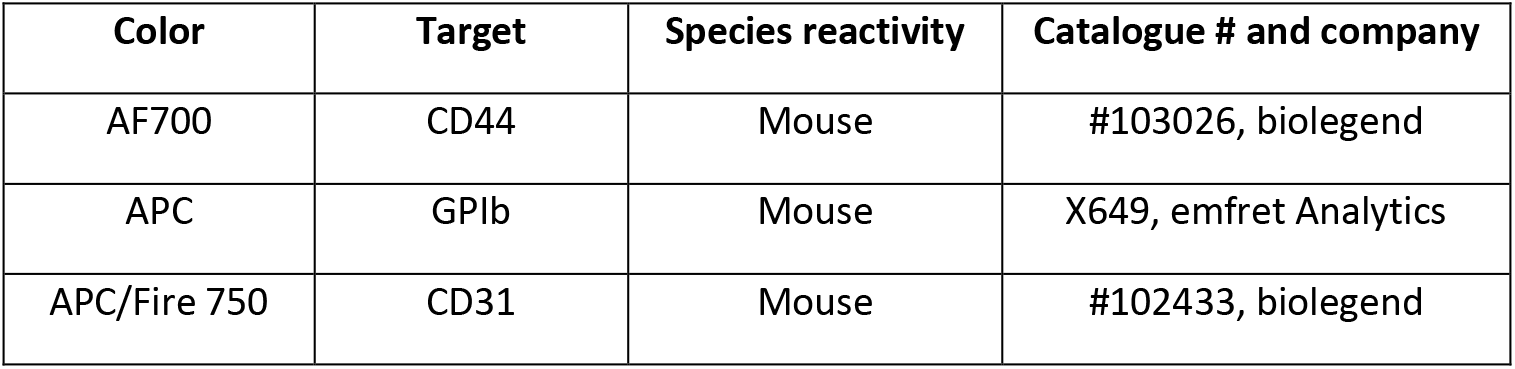
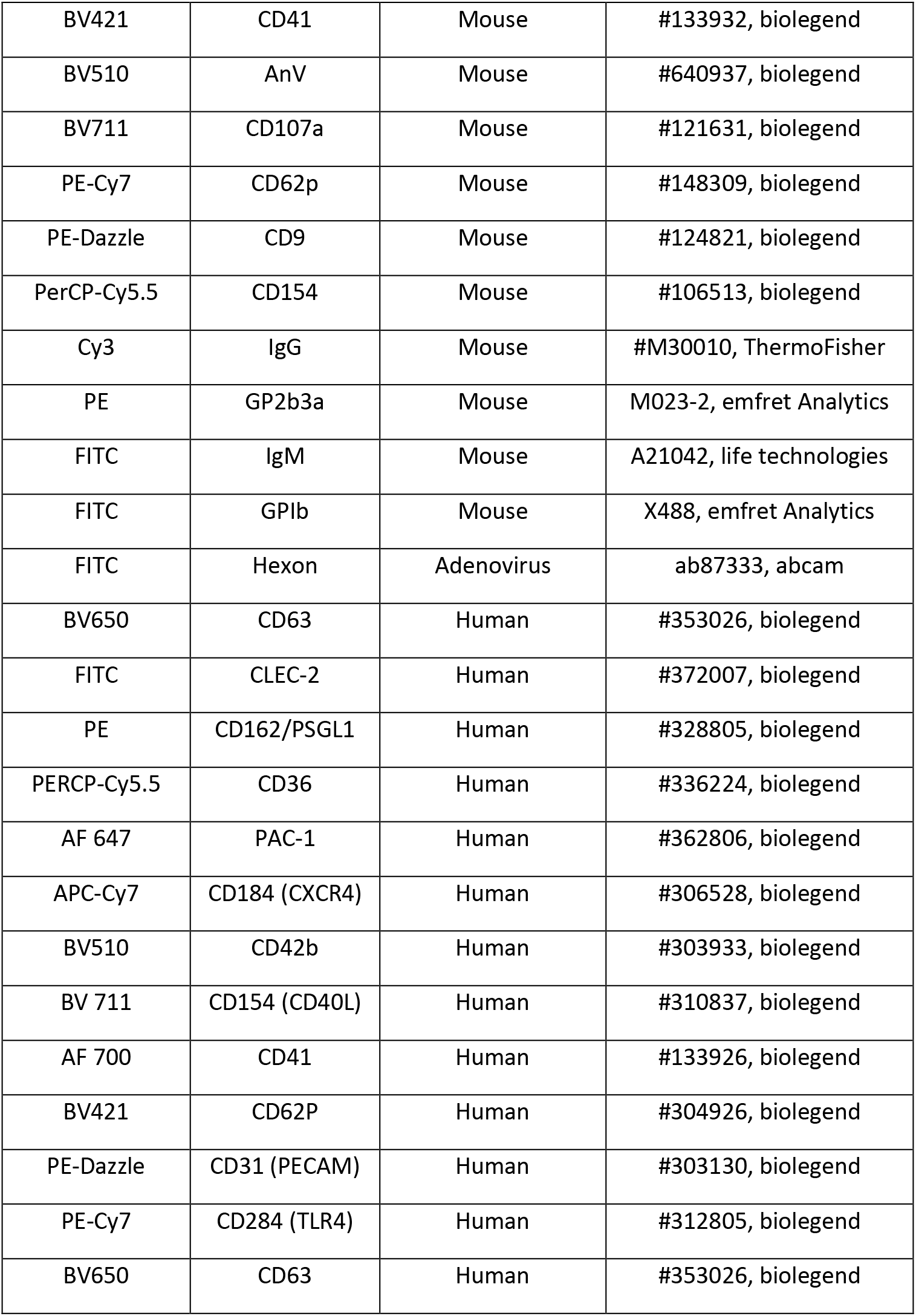

### Histology

Organs were harvested directly after sacrifice, fixed at 4%PFA for 1h and subsequently 30% sucrose overnight, then embedded in OTC and stored at −80°C. Sections were stained with antibodies against F4/80 (Biolegend, Cat. No. 123110), CD45R/B220 (Biolegend, Cat. No. 103212), ki67 (Abcam, Cat. No. ab15580), Lectin PNA (Thermofisher, Cat. No. L32458). Secondary antibodies and nucleic acid stain included FITC goat anti mouse (Thermofisher, Cat. No. A11029) and Cy3 goat anti rabbit (Jackson Immunoresearch, Cat. No. 111-165-003) and Hoechst33342. Micrographs were taken on a Zeiss LSM 880 confocal microscope in airyscan mode and a Leica LAS X epifluorescence microscope.

### Data analysis and statistics

Analysis of histology and intravital imaging was done using ImageJ v2.1 or Imaris (Bitplane). Intravital imaging cell tracking was done manually on randomly selected cells which could be traced for at least 5 frames. Statistics were computed with Imaris, Meandering Index was defined as Net displacement/Total track length. Unpaired t-tests with Welch’s correction was used for all cell tracking analyses. For area quantification, thresholds were taken with ImageJ and overlap quantified by percent area overlapping between thresholds. For all statistical tests Prism (GraphPad) was used. Unpaired t-test were used unless otherwise noted. All data are shown as mean ± standard error of the mean (s.e.m.). When more than three t-tests were applied on the same data set, multiple testing correction was done.

**Supplementary Figure 1.**
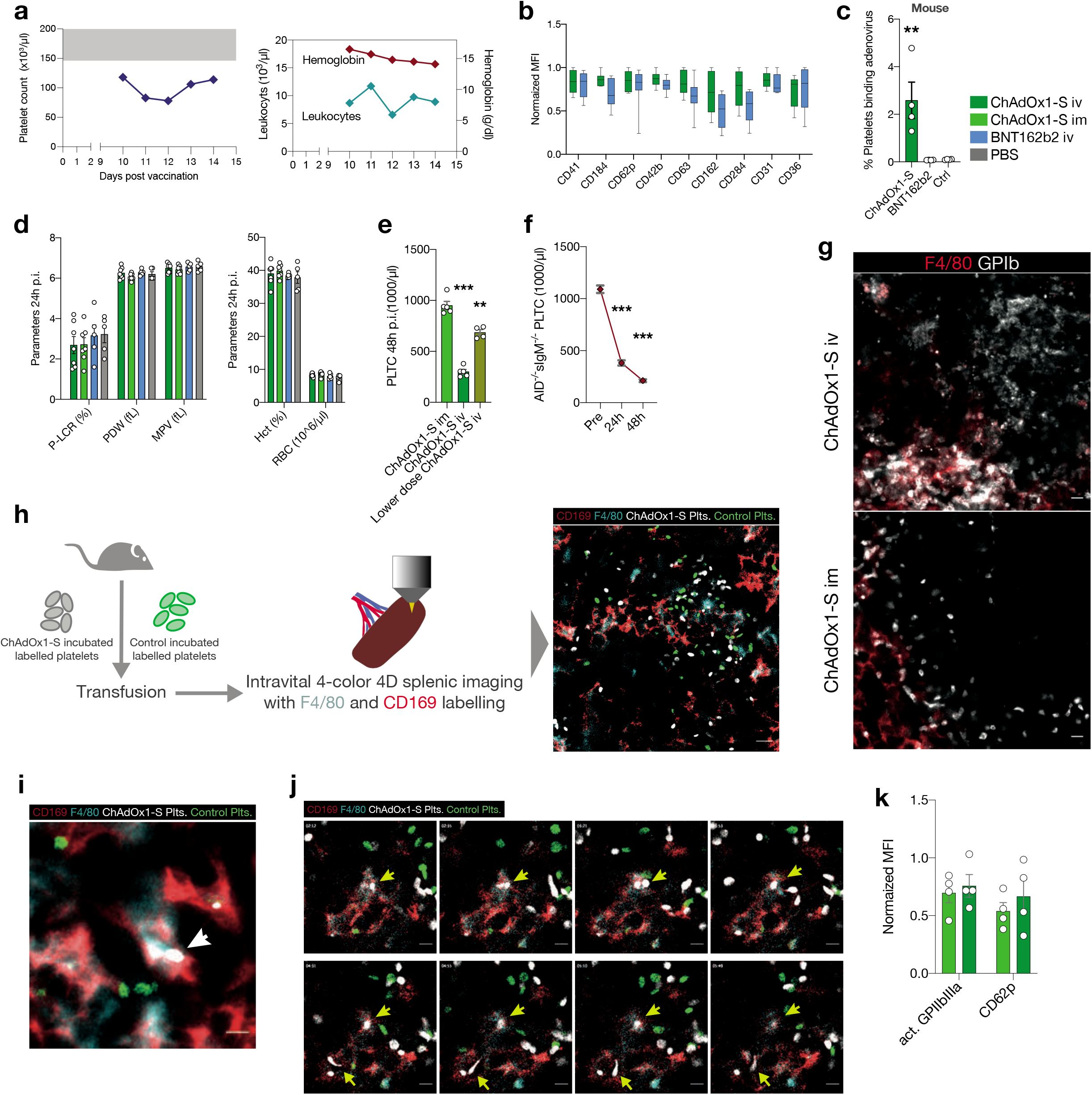
**a,** Time course of hemoglobin, leukocyte and platelet counts of SVT patient. **b,** Platelet surface marker expression of human platelets incubated with ChAdOx1 nCov-19 or BNT162b2. Normalized MFIs. Multiple t-tests with Holm-Sidak correction, non-significant. n=8 donors per group. **c,** Quantification of adenovirus platelet binding to mouse platelets. One-way ANOVA with post-hoc Tukey’s test. Comparison of ChAdOx1 nCov-19 to both controls. n=4 per group. **d,** Cell count parameters of blood taken at 24h p.i. with either ChAdOx1 nCov-19 i.v. or i.m., BNT162b2 i.v. or PBS i.m.. Two-way ANOVA with post-hoc Tukey’s test, all nonsignificant. n≥5 per group. **e,** Comparison of platelet count at 24h p.i. with either ChAdOx1 nCov-19 i.v. or i.m. (same data as in Figure 1f) or low-dose ChAdOx1 nCov-19 i.v. administration. Unpaired t-tests, n≥4 per group. **f,** Time course of platelet counts right before and after ChAdOx1 nCov-19 i.v. administration of AID^-/-^sIgM^-/-^ mice. Unpaired t-tests, n=4 per time point. **g,** Zoom-in of the crop outs in Figure 2a. Scale bars are 5μm. **h,** Illustration of intravital splenic imaging with transfused platelets and representative overview image of intravital microscopy. Scale bar 10μm. **i,** Z-projection of a 3D-stack intravital splenic microscopic image showing phagocytosed ChAdOx1 nCov-19 pretreated transfused platelets (white, arrow) in comparison to PBS treated platelets (green). Scale bar 5μm. **j,** Series of images from an intravital splenic microscopic video showing phagocytosis (upper arrow) and interactions (lower arrow) of ChAdOx1 nCov-19 pretreated transfused platelets. Time is shown on the upper left, scale bars 5μm. **k,** Platelet activation marker expression after plasma incubation of mice with i.v. or i.m. ChAdOx1 nCov-19 administration. Normalized MFIs. Unpaired t-tests, non-significant. n=4 mice per group. Error bars are mean ±s.e.m. *p<0.05, **p<0.01, ***p<0.001.

**Supplementary Figure 2.**
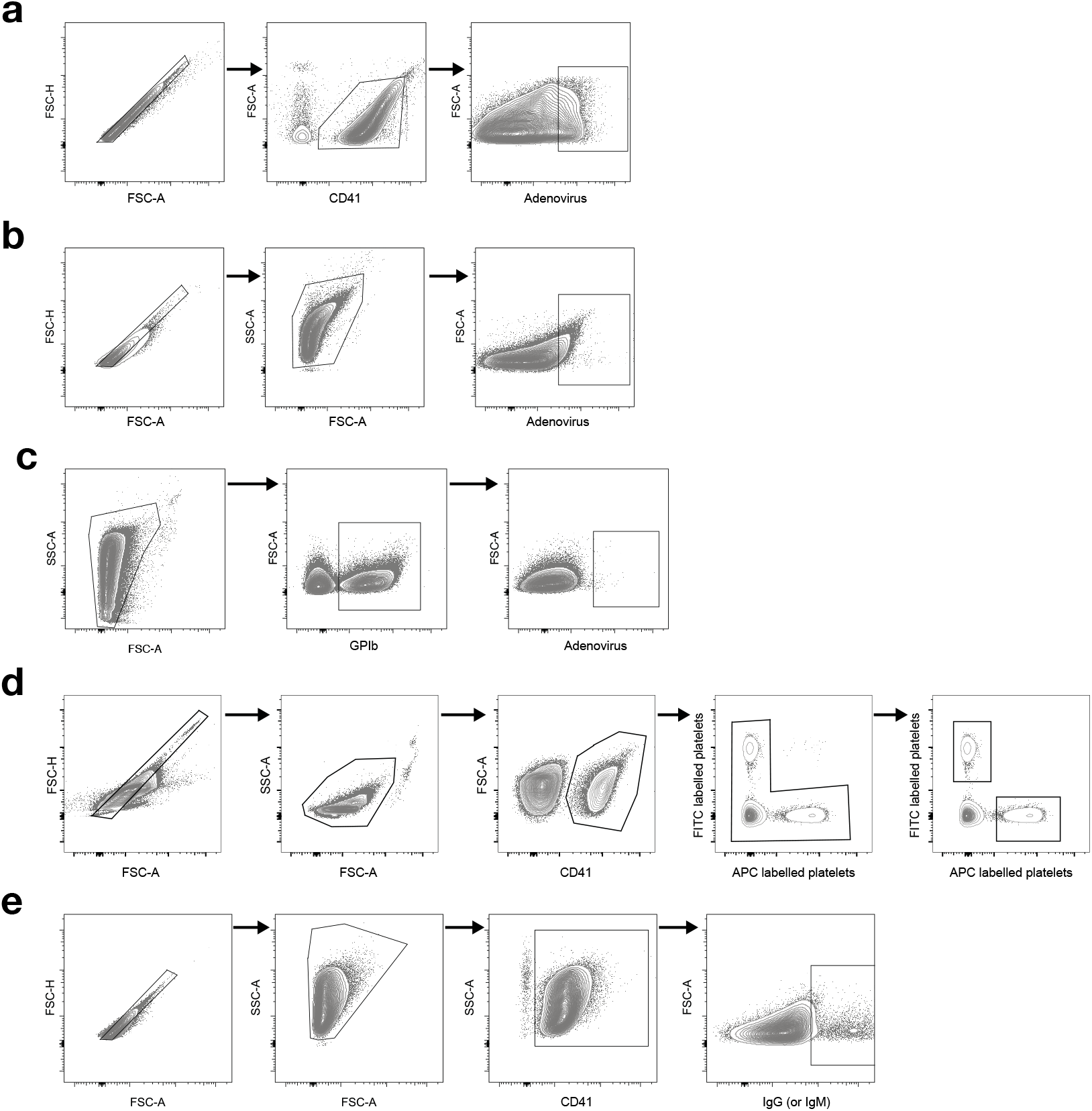
**a,** Gating strategy for human in-vitro platelet-adenovirus binding. Last gate was used to quantify platelet-adenovirus binding. **b,** Gating strategy for murine in-vitro platelet-adenovirus binding. Last gate was used to quantify platelet-adenovirus binding. **c,** Gating strategy for murine in-vivo platelet-adenovirus binding and platelet surface marker expression. Second gate was used to derive MFIs for platelet surface marker expression, last gate was used to quantify platelet-adenovirus binding. **d,** Gating strategy for transfused platelet tracking. Both of the last gates were used to quantify ChAdOx1 nCov-19 and BNT162b2 incubated transfused platelet fraction. **e,** Gating strategy for immunoglobulin binding of platelets incubated with plasma from vaccinated mice. Last gate was used to quantify either IgG (shown) or IgM binding. All gating is shown as contour plots with 2% counter and outliers shown.

**Supplementary Figure 3.**
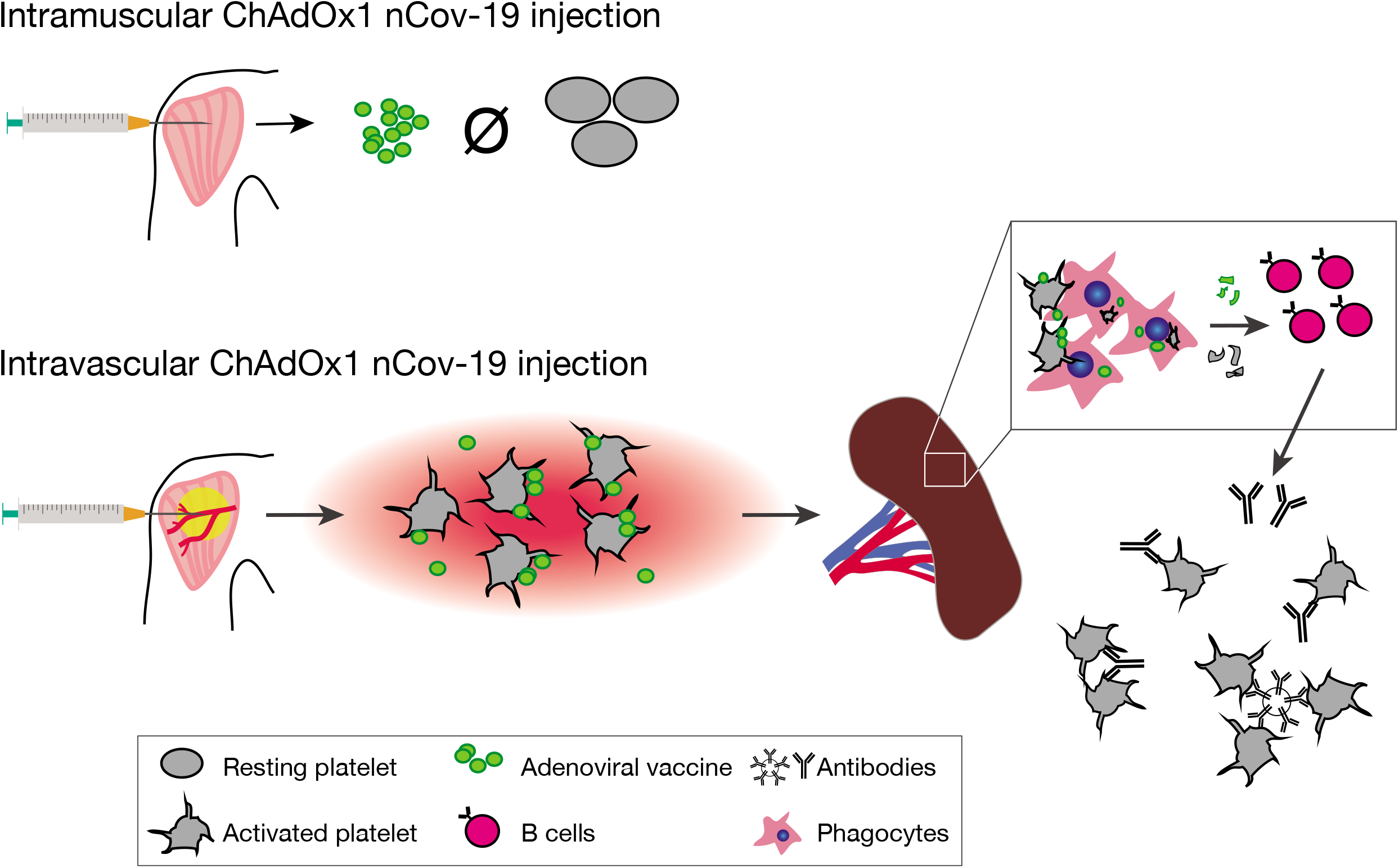
Graphical abstract: Thrombocytopenia and splenic platelet directed immune responses after intravenous ChAdOx1 nCov-19 administration. Intramuscular injection of ChAdOx1 nCov-19 leads to normal vaccine response without platelet involvement. Accidental intravascular injection of ChAdOx1 nCov-19 leads to adenovirus-platelet binding and platelet activation. Platelets are cleared by professional phagocytes, particularly in the spleen. Trafficking and processing of platelet adenovirus-aggregates from the red pulp to splenic follicles leads to a B cell response with the emergence of platelet binding antibodies.

## Author contributions

LN and AL conceived of and initiated the study. LN conceptualized the study. LN, AL, KP, KS and SM planned the experiments. LN, AL, KP, AA, ER, MH, VP, LE, DR and AT conducted the experiments. AL, LN, KP, AA, RE and RK analyzed the results, AL visualized the findings. LN wrote the initial draft, LN, AL, KP, KS and MS edited the manuscript with input from all authors. KaS and MI provided resources. LN, KP, KS and SM provided oversight, resources and funding.

We thank all lab members for their valuable input and help, as well as Dr. Schneider for his substantial support.

## Materials & correspondence

Materials are shared upon reasonable request by email.

## Corresponding authors

&, Medizinische Klinik und Poliklinik I, University Hospital Ludwig-Maximilian-University Munich, Marchioninistr. 15 81377, Munich, Germany.

