## Supplementary material for "Thrombocytopenia and splenic platelet directed immune responses after intravenous ChAdOx1 nCov-19 administration": Suppl. Material and Methods

### Suppl. Methods section

#### Patient data

The patient was treated at the university hospital LMU Munich. The patient consented to taking part in this study. The study was approved by local ethics committee of Munich. Data on presentation, treatment and outcome are depicted in Fig 1a-b, Suppl. Fig 1a and the laboratory values table (Suppl Table 1-3). A HIT-IL-Acustar assay and a HYPHEN BioMed HIT-ELISA Test deemed sensitive for TTS returned negative for this patient [5]. PF4 induced platelet activation(PIPA) and heparin induced platelet activation (HIPA) tests were performed as previously reported[24].

Suppl. Table 1

| **Laboratory values at admission** | | | |
|---|---|---|---|
| Parameter | | Unit | Range |
| INR | 1.1 | - | 0.8-1.2 |
| aPTT | 31 | sec | 22-34 |
| CRP | 0.2 | mg/dl | ≤0.5 |
| D-Dimer | 1.6 | µg/ml | ≤0.5 |
| Leukocytes | 8.68 | G/l | 3.9-9.8 |
| Hemoglobin | 16.5 | g/dl | 13.5-17.5 |
| Platelets | 118 | G/l | 146-328 |
| **Thrombophilia screening** | | | |
| Factor-V-Leiden mutation | 1691G>A not detected | | |
| Prothrombin mutation | 20210G>A not detected | | |
| Lupus anticoagulant | not detected | | |
| APC screening | 4,4 (range >=2,5) | | |
| Protein C | 78% (range 68-150) | | |
| Protein S | 91% (range 74-136) | | |

Suppl. Table 2

| **HIT tests** | |
|---|---|
| Heparin/PF4 Ab Screening Test | **positive** |
| HIT-IL-Acustar-Assay | negative |
| HIT-ELISA-Test (HYPHEN Biomed) | negative |
| HIPA test | negative |
| PIPA test | negative |

Suppl. Table 3

| **Platelet auto-antibody panel** | |
|---|---|
| PakAuto® Test* | **positive** |
| GPIb/IX Auto-antibodies* | negative |

Antibodies used for Flow Cytometry:

| Color | Target | Species reactivity | Catalogue # and company |
|---|---|---|---|
| AF700 | CD44 | Mouse | #103026, biolegend |
| APC | GPIb | Mouse | X649, emfret Analytics |
| APC/Fire 750 | CD31 | Mouse | #102433, biolegend |

| | | | |
|---|---|---|---|
| BV421 | CD41 | Mouse | #133932, biolegend |
| BV510 | AnV | Mouse | #640937, biolegend |
| BV711 | CD107a | Mouse | #121631, biolegend |
| PE-Cy7 | CD62p | Mouse | #148309, biolegend |
| PE-Dazzle | CD9 | Mouse | #124821, biolegend |
| PerCP-Cy5.5 | CD154 | Mouse | #106513, biolegend |
| Cy3 | IgG | Mouse | #M30010, ThermoFisher |
| PE | GP2b3a | Mouse | M023-2, emfret Analytics |
| FITC | IgM | Mouse | A21042, life technologies |
| FITC | GPIb | Mouse | X488, emfret Analytics |
| FITC | Hexon | Adenovirus | ab87333, abcam |
| BV650 | CD63 | Human | #353026, biolegend |
| FITC | CLEC-2 | Human | #372007, biolegend |
| PE | CD162/PSGL1 | Human | #328805, biolegend |
| PERCP-Cy5.5 | CD36 | Human | #336224, biolegend |
| AF 647 | PAC-1 | Human | #362806, biolegend |
| APC-Cy7 | CD184 (CXCR4) | Human | #306528, biolegend |
| BV510 | CD42b | Human | #303933, biolegend |
| BV 711 | CD154 (CD40L) | Human | #310837, biolegend |
| AF 700 | CD41 | Human | #133926, biolegend |
| BV421 | CD62P | Human | #304926, biolegend |
| PE-Dazzle | CD31 (PECAM) | Human | #303130, biolegend |
| PE-Cy7 | CD284 (TLR4) | Human | #312805, biolegend |
| BV650 | CD63 | Human | #353026, biolegend |
